# FishProx: Proximate conspecific interaction of Atlantic salmon (*Salmo salar*) for behavioural analysis through instance segmentation masks

**DOI:** 10.1101/2025.07.07.659286

**Authors:** I-Hao Chen, Nabil Belbachir, Lars Ebbesson, Antonella Zanna Munthe-Kaas, Lars Erik Solberg, Gaute Alexander Nedberg Helberg, David Izquierdo-Gomez, Santhosh K. Kumaran, Jelena Kolarevic, Chris Noble

## Abstract

In modern fish production and experimental facilities, camera installations are increasingly common for monitoring animal welfare. This opens up a new alley of autonomous tracking systems levering streamed data. However, the substantial amount of data necessitates condensation for human operators. In this paper, we present the out-of-the-box pipeline called FishProx that starts with the imagery of fish in tanks and ends in the calculation of welfare impacting factors through anomaly scores. The utilisation of the Segment Anything Model (SAM) is the basis for the identification of feeding behaviour and also (ab)normal behaviour through analysing snapshots of the fish tanks to infer metrics such as fish cohesion and cluster alignment, alongside the detection count. Our method employs automatic and robust algorithms for camera systems, making additional labelled data unnecessary, thereby saving energy and costs. Furthermore, automatic fish observation coupled with integrated analytical capabilities can lead to a sensible autonomous decision-making process. Lastly, by providing objective metrics, our approach might guide the enabling of automatic quantification of fish states, making comparative studies possible.

## 1. Introduction

“Fish that are close to each other, should swim together”. This is the idea leading to the pipeline FishProx, which implements a microservice-oriented architecture to utilise the Segment Anything Model (SAM) [9] to facilitate downstream tasks like local fish behaviour analysis.

**Figure 1.**
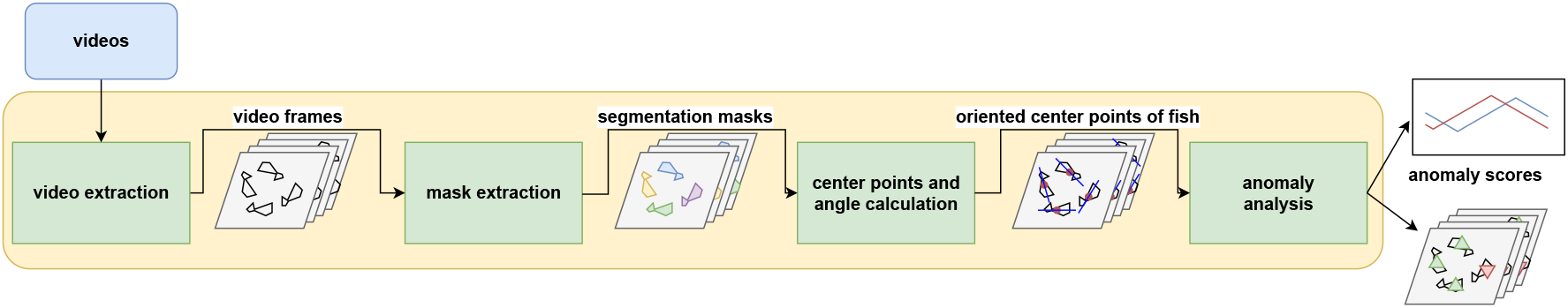
Schematics of FishProx: The segment anything model (SAM) [9] is used on extracted frames to prompt segmentation masks that are filtered for fish segmentations. Afterwards, the segmentation centers and their 180*^°^*-agnostic orientation are calculated. To obtain anomaly scores, segementation centers are clustered with K-means (K=16) which gives rise to the fish cohesion score (intra-cluster) and the cluster alignment score (inter-cluster) using the in-and outliers of the orientation angles. Integrating these scores with the segmentation detection count enables the quantification of some fish behaviour in RAS tanks.

### 1.1 Related Work

In the past few years, generative AI is hitting our mainstream society around the world [4, 13, 22]. Natural language processing (NLP) [5], generative pre-trained transformers (GPT) [19, 26] and large language models (LLM)[18] have gained in maturity over the past years to show impressive capabilities in the everyday life as tools ranging from education [3, 15, 16] to content creation [24].

At the same time, generative models for animal behaviour are not widespread. The labelling and tracking of animals [14] for various purposes have been explored, though not for fish. Work on zero-shot detection for species recognition can be found [21], but to the best of our knowledge, there exists no integrated pipeline or model that generates meaningful metrics for fish populations from video data.

Work on fish segmentation itself, on the other hand, has been studied using various architectures [7, 10, 23, 25], and with the rise of the Segment Anything Model (SAM) [9], the way segmentation can be obtained was also revolutionised for the aquatic sector.

### 1.2. Recirculating aquaculture system (RAS)

Recirculating aquaculture systems (RAS) have been under heavy development in the past two decades, which lead to increased size and fish capacity and optimized resource usage [6]. Using RAS to farm fish has advantages in the environment-friendly and sustainable area [11]. It is water efficient and highly productive and is often protected from environmental impacts such as pollution and parasitic infections [2]. Current trends suggest an increase of RAS facilities around the world inline with an increase in production of Atlantic salmon *Salmo salar*), one of the major farmed fish species [1].

### 1.3. Experiment Details

The RAS research facility used in this experiment was located in Sunndalsøra, Norway. Each of the six water tanks had a volume of 3.3*m*^3^ and was shaped like an octagon. Additionally, a top camera, a IR Dome (Imenco AS) was installed above each tank with a video resolution of 1080p @ 30fps. Another camera was placed underwater, and a hydrophone (the long rod through the pictures in Figure 2) was installed for each tank. A trial experiment was ongoing for several months in 2023 (May until August), which is not the focus of this paper. A total of 1000 Atlantic salmon were in each tank with a weight of 40 grams ± 10 grams at the starting date of 31.03.2023. The data available for this study was recorded at tank 210 and consists of around 8 hours of video material from 12.05.2023.

**Figure 2.**
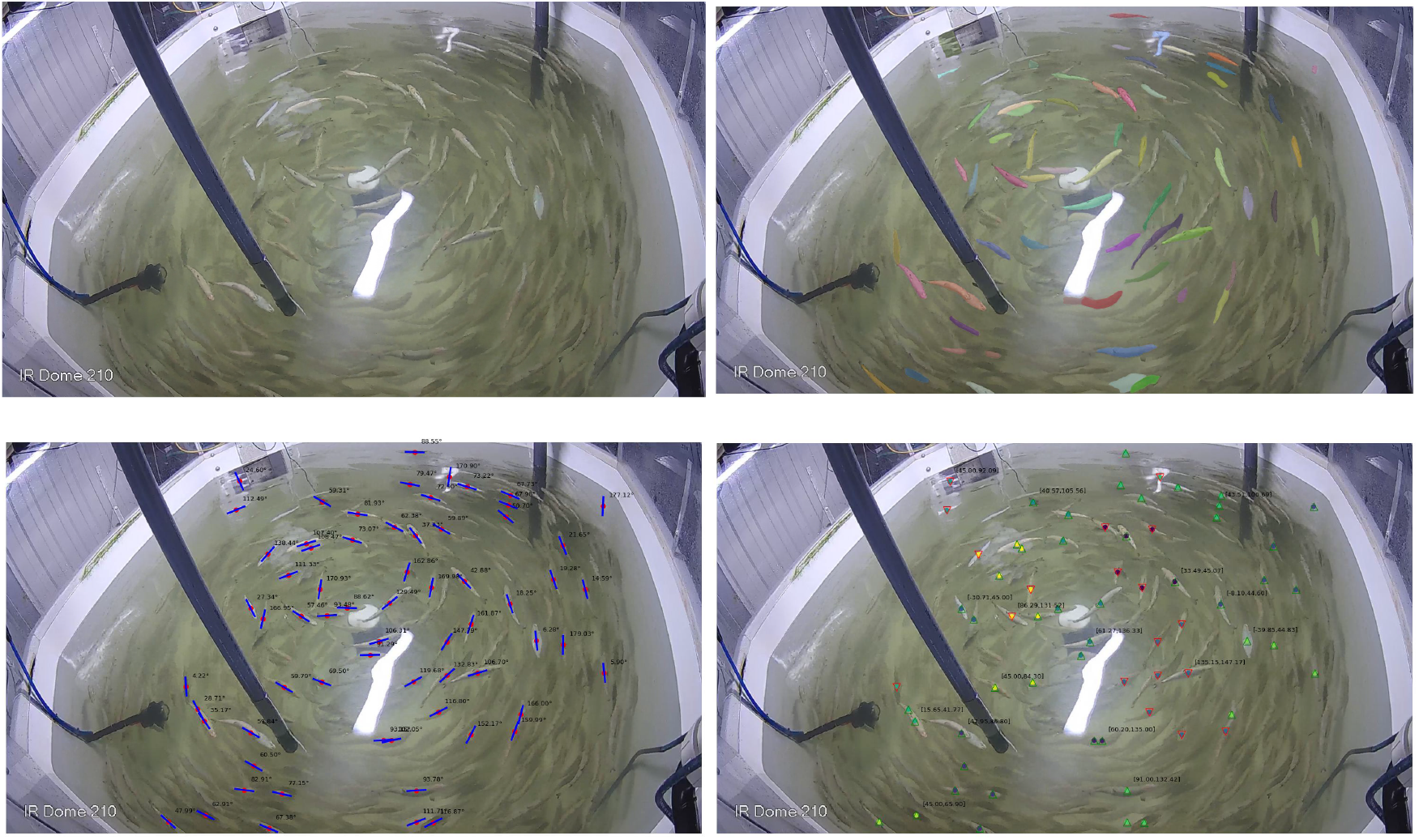
In sequential order: Original image of the fish RAS tank 210, the tank with fish segmentation based on SAM [9], then orientation angles are displayed for each fish segmentation, and lastly, the classification that leads to anomaly scores (see Section 2.4.1).

## 2. The FishProx pipeline

The FishProx pipeline consists of multiple Python code modules that were put in docker images [17] and orchestrated by a dockerfile. The running services were communicating through the file-system which allowed for manual inspection. Chained together, they formed a robust video processing pipeline which could be used OS independently. We present the modules in order:

### 2.1. Video Extraction

While this service was running, new videos were just dropped into its overseen folder, triggering the processing into frames in a parallel manner.

### 2.2. Frame Analysis

Frames added to the input folder of this service triggered SAM [9] (checkpoint sam vit l 0b3195) to prompt segmentation masks for the frames. This was programmed concurrently with shared GPU such that we achieve full utilisation of the GPU (Nvidia GeForce RTX4090) as this step was usually the bottleneck. The segmentation result for each frame from SAM was saved into the file system as a .json file.

### 2.3. Mask Processing

#### 2.3.1 Mask Selection

From all the segmentations of the fish tank, only fish segmentations were of interest. A filtering technique based on Kernel Density Estimation (KDE) [20] was applied to the pixel size of the segmentations in conjunction with creating KDE “groups” through thresholding with local extrema. Here we are leveraging knowledge about the fish size of the same rearing process which have similar sizes and therefore being grouped into the same KDE “group”. Additionally, the convex hull of the tank segmentation was calculated to account for the other objects occluding the water surface and then used as a condition for the fish masks.

The center of each fish segmentation was calculated by taking the center of mass of the segmentation. Additionally, each fish segmentation was tagged with the water tank section it appeared in. The tank sections ranged from 1 to 8 and were evenly anti-clockwise distributed around the centrally located water outlet in the RAS tank. Section One was marked by the 45^°^ slice by taking the right horizontal from the water outlet and rotating it anti-clockwise. Outputs of this service were the center points of each detected fish individual and the finished segmented masks for each frame which gave rise to the first metric: The number of detected fish in the frame, alias the detection count score.

#### 2.3.2 Calculation of Orientation Angles

PCA [8] was used to obtain the main axis of fish for orientation. The biggest Eigenvector (*x, y*) was used to calculate the orientation angle:

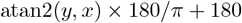

The resulting angles were in the interval of [0^°^,360^°^). We did not differentiate between forward or backward movements, since we considered both swimming with and against the water currents valid swimming directions. Therefore, we applied the surjection mod _180_ : → [0, 360) → [0, 180) with *x* → *x* mod 180 to overlap the angles are 180^°^ apart. This resulted in a 180^°^ agnostic orientation and the angle value domain was modelled by the interval [0^°^, 180^°^). A fish that had the orientation of a vertical line (facing upwards or downwards) thereby had an angle of 0^°^, and a fish with horizontal alignment had an angle 90^°^.

### 2.4. Checking Proximate Swimming Behaviour

We clustered the fish with K-means [12] with *K* = 16 and for each cluster *k* ∈ {1*, …,* 16}, the family of *N_k_* orientation angles 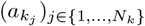 were taken to calculate the circular MAD (Median Absolute Distance). To preserve the 180^°^ periodicity, we normalised the orientation angles to radians with the scaling factor 2*π/*180 to calculate their position on the unit circle with sine and cosine. Then, the median of the sines and cosines was calculated. We note that we made use of the notation 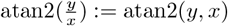. For each cluster group *k* we calculated:

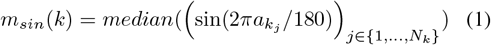

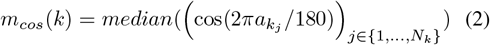

and then re-transforming the median values back into an angle

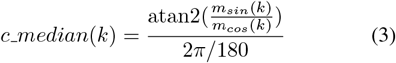

Lastly, we add 180 if the *c median*(*k*) is negative to push all values into the positives.

For each cluster element in cluster *k*, the median absolute distance was calculated component-wise using *m_sin_*(*k*) and *m_cos_*(*k*) as medians and calculating in the distance after transforming the angle of the cluster element into a point on the unit circle:

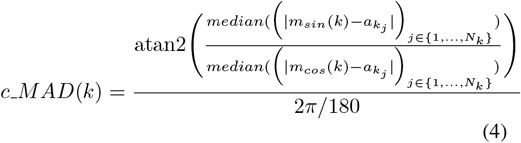

Fish whose orientation angles fell outside of the interval *c normal*(*k*) := [*c median*(*k*) − *c MAD*(*k*), *c median*(*k*) + *c MAD*(*k*)] were marked as *anomalous* (red) for swimming different from their close peers. Otherwise, fish that swim uniformly by the same criteria were marked *normal* (green) (see Figure 2).

#### 2.4.1 Anomaly Scores

The fish cohesion score was calculated by dividing the number of normal swimming fish by the total number of fish segmentations. The cluster alignment was calculated by dividing the number of K-means clusters with zero anomalous segmentation centers by the number of all clusters.

## 3. Results

The results were aggregated in one-minute intervals.

### 3.1. Feeding Behaviour

Automatic feeding of fish started at 9:00 and ended at 11:00 on the 12th of May 2023, as marked in green in Figure 3 and 4. There was only a slight increase in the fish cohesion at the start of the feeding time, but cluster alignment spiked up at the feeding start at 9:00 and persisted for half an hour before normalising itself after an hour. The number of detections dropped sharply to around half of their previous level in Figure 4 but also recovered within the next hour. Especially the sections five to eight which lied close to the feeder position display a heavy drop in counts. Note that the metrics of fish cohesion and cluster alignment are inverse to the detection count.

**Figure 3.**
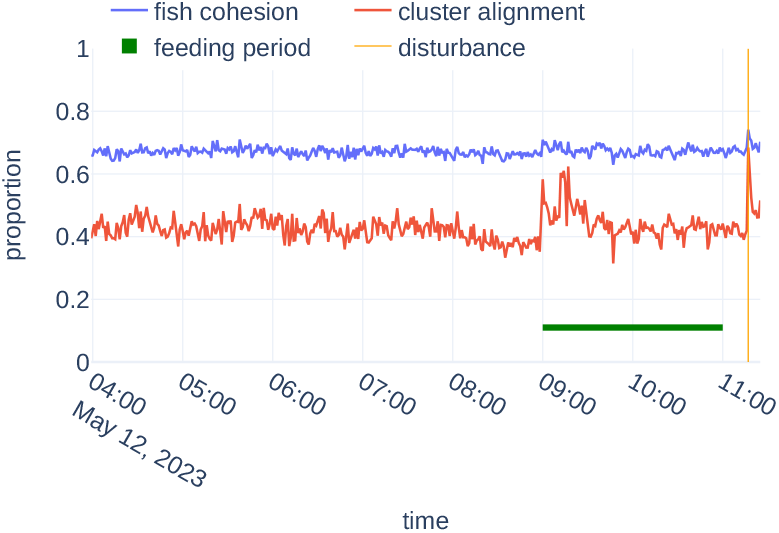
Fish cohesion and cluster alignment as anomaly scores. The green horizontal line indicates the feeding period, and the vertical orange line the tank cleaning event at 11:17.

**Figure 4.**
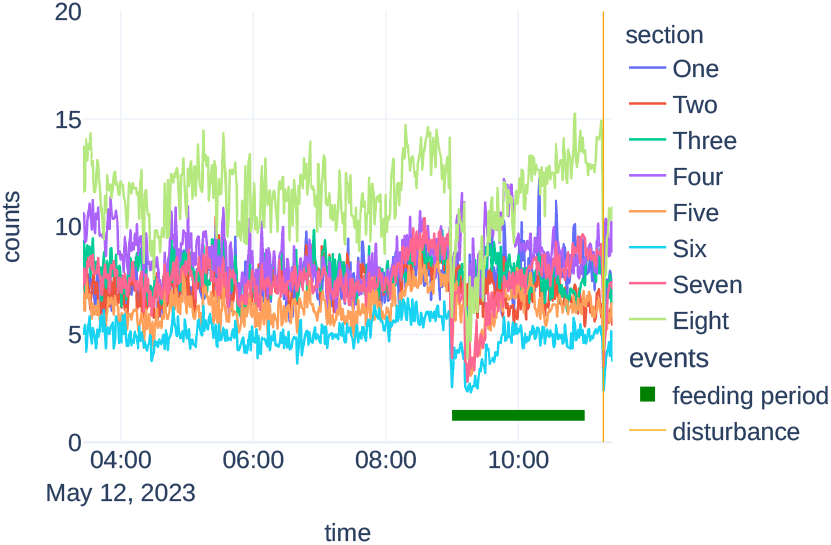
Detection counts by tank section. The green horizontal line indicates the feeding period, and the vertical orange line the tank cleaning event at 11:17.

### 3.2. Disturbance Event

One special datapoint of interest was 11:17. A disturbance to the fish group was captured on video when the fish population rapidly evacuated one area of the tank due to the in tank cleaning operations carried out by a member of staff. This behavioural response can be indicative of exposure to a stressor. This single event was also visible in Figure 3, being the most prominent spike in both fish cohesion and cluster alignment.

## 4. Discussion

Fish segmentation is a difficult task, but with stable conditions as in RAS, we can analyse video streams efficiently. It also allows to use of strong reflection and unresting water as a source of information, leading to the metric of counting the number of fish detections. We were able to show that both feeding and tank cleaning events could be measured with the presented metrics. We also showed that the disturbance due to tank cleaning leads to a higher spike, meaning a stronger reaction of fish in the tank was captured accordingly by the pipeline. The inverse relationship between detection counts and the anomaly scores is expected, since when the number of detections in total drops, and we assume an even distribution of cluster centers by K-means, then each cluster will have fewer fish individuals, making it more likely to be in the orientation angle interval in Section 2.4. The main drawback of this pipeline is the horizontal scalability and reliance on stable videos. Bigger tank setups may require different camera positioning or it may not be feasible to have the whole tank in scope.

## 5. Conclusion

In this study, we presented the deployable pipeline FishProx for fish state observation. We hope to push further the digitisation of the aquaculture industry with the presented readable metrics by providing quantification of events such as feeding and disturbances. Insights on anomalous behaviour in fish held in RAS can be useful to a more holistic view on fish welfare which is one of the steps towards an automated welfare assessment system. Further work would encompass the implementation of real-time sensors with pipelines like FishProx to make datastreams easily accessible and interpretable for facility personnel.

## 6. Funding

This work was funded by the RAS4.0 project through Research Council of Norway (RCN) under the project number 320717 and the project grant with number 323300. The authors would like to thank Britt Kristin Megård Reiten for the technical expertise and assistance during the study.

## Notes

### Competing Interest Statement

The authors have declared no competing interest.

